# Experimental evidence that UV/yellow colouration functions as a signal of parental quality in the blue tit

**DOI:** 10.1101/2020.09.14.293613

**Authors:** Jorge García-Campa, Wendt Müller, Judith Morales

## Abstract

In bi-parental species, reproduction is not only a crucial life-history stage where individuals must take fitness-relevant decisions, but these decisions also need to be adjusted to the behavioural strategies of a partner. Hence, communication is required, which could be facilitated by condition-dependent signals of parental quality. Yet, these traits have (co-)evolved in multiple contexts within the family, as during reproduction different family members may coincide and interact at the site of breeding. In this study we explore whether a condition-dependent trait acts a quality signal and regulates intra-family interactions in a bird species, the blue tit (*Cyanistes caeruleus*). As a family is a complex network where signals could be perceived by multiple receivers, we expected that experimentally blocking the reflectance of an adult’s UV/yellow colouration of breast feathers may affect the behavioural strategies of all family members. We found an increase of parental investment in nests with an UV-blocked adult, as the partner compensated for the perceived lower rearing capacity. As the UV-blocked adult did not change its provisioning behaviour, as was to be expected, their partner must have responded to the (manipulated) signal but not to a behavioural change. However, offspring did not co-adjust their begging intensity to a signal of parental quality. Opposite to adults, we propose that offspring respond to the behaviour but not to the parental signal. Overall, our results show experimentally at the first time that UV/yellow colouration of blue tits acts as a quality signal revealing the rearing capacity to mates.

## Introduction

Communication is a co-evolutionary process that involves the transfer of information from a transmitter to a receptor individual (Bradbury & Vehrencamp 1998). This process is subject to selection, where both transmitters and receivers try to optimize their fitness as a result of their interaction (Zahavi 1981). Communication usually involves conspicuous traits (i.e., colourations or vocalizations) that can be costly to produce and maintain (Zahavi 1975; Hamilton & Zuk 1982). Therefore, these traits typically reflect health or condition (Folstad & Karter 1992), and may thus function as signals of quality. Individuals respond to these signalling traits to modulate fitness-related decisions - like where to live, what to eat and whom to interact with (Danchin et al. 2004).

One important time point where fitness-related decisions require communication is during reproduction, when, at least in bi-parental species, individuals must (co-)adjust the amount of parental care they are going to provide (Iserbyt et al. 2019). Because parental investment in current reproduction comes at a cost to parents in terms of survival and thus future reproduction (Stearns 1992), each parent would benefit from shifting the offspring investment to their mate (Trivers 1972). Furthermore, to enhance reproductive success, caregivers must adjust their level of investment according to the quality of the current reproductive event, which depends on i) the environmental conditions like habitat quality (Capilla-Lasheras et al. 2017) or resource availability (Hakkarainen et al. 1997), ii) their own condition, which determines their rearing capacity (Ots & Horak 1998), and iii) the condition of their mate, which can be reflected by signals of quality (Velando et al. 2006).

How individuals respond to signals of genetic or phenotypic quality has received significant attention (Sheldon 2000; Harris & Uller 2009; Ratikainen & Kokko 2010). On the one hand, individuals are expected to increase their effort when paired with highly ornamented mates either because the offspring will inherit the mate’s high quality conveyed by ornamentation (i.e., the classical scenario of positive differential allocation; Burley 1986; 1988), or because highly ornamented mates seek more extra-pair copulations and thus neglect their parental effort (Kokko 1998; Ratikainen & Kokko 2010). On the other hand, individuals are also expected to increase their investment when paired with poorly ornamented mates (or in general poor quality mates) in order to improve on an unfortunate situation, especially if the later occurred unexpectedly or is expected to be time-limited (reproductive compensation; Gowaty et al. 2007; 2008). Whether an increase or decrease in parental effort should be expected is thus context-dependent, but as this is about communication, it is also highly dependent on the behaviour of the signaller (Ratikainen & Kokko 2010). However, previous studies have primarily focused on the receiver’s response, and have not always controlled for the signallers’ behaviour and its investment in reproduction (but see, for example, Sanz 2001; Soler et al. 2008; Doutrelant et al. 2008), while this is crucial to understand the occurrence of differential allocation and reproductive compensation.

Additionally, studies have generally focused on a single set of receivers, for example, the focus has been on the response of parents to signals expressed by mates, but not on offspring responses to parental signals. However, family members (mates and offspring) usually coincide in time and space, and the information conveyed by signallers can be used simultaneously by multiple family members. Hence, the family is a social environment in which signals can function in multiple contexts (Morales & Velando 2013), being possibly perceived by offspring (Lucass et al. 2016) and even helpers in species with cooperative breeding (Canestrari et al. 2011). To take an example, the size of the red-dot on the bill of the yellow-legged gull, *Larus michaellis*, affects the mate’s parental investment (Morales et al. 2009) and at the same time it induces chick begging (Velando et al. 2013). Similarly, in the burying beetle, *Nicrophorus vespilloides*, chemical profiles play a role in mate-mate recognition (Steiger et al. 2008) and also triggers larvae begging (Smiseth et al. 2010). Studies investigating signals expressed in the family environment should hence assess their effect(s) on multiple receivers.

Here, we studied whether an adult trait that reflects condition (i.e., UV-based plumage colouration) functions as a signalling trait that affects the mate’s parental effort, while controlling for the signaller’s own effort. Moreover, given that the family is a social environment, we studied whether the trait affects the offspring decisions as well. To test these hypotheses, we experimentally blocked the reflectance of the UV region of yellow breast feathers of blue tits (*Cyanistes caeruleus*), a socially monogamous passerine with bi-parental care, which represents an ideal study model of parental investment and signalling. UV-based colouration in different structures has been found to signal quality or condition (White 2020) and to play a role in mate choice as well as in parental decisions after mating (Hunt et al. 1999; Limbourg et al. 2013a). UV-based colouration expressed by offspring is also known to affect parental decisions (Jourdie et al. 2004; Morales & Velando 2018). We manipulated one of the parents in each experimental nest to simulate low quality. We then explored the parental effort of the manipulated individual and that of its mate, as well as chick begging behaviour. We predict that parents will increase their effort when mated with UV-blocked individuals in order to compensate on a low-quality situation, given that apparently their mate has a reduced condition and thus low rearing capacity. However, we expect that UV-blocked individuals do not change their own parental effort, given that their inherent quality has not truly changed but only their external appearance (i.e., signal expression), which should be perceived by mates as reduced condition. We also expect that offspring could perceive UV-blocked parents as poor-quality caregivers, and thus should redirect their begging behaviour towards non-manipulated parents.

## Material and methods

### Ethics statement

According to the Spanish laws in relation to animal research, the study licenses to perform the study were approved by the Spanish Research Council (CSIC, ref. (ref. 639/2017) and the Consejería de Medio Ambiente, Administración Local y Ordenación del Territorio, Comunidad de Madrid (ref. 10/056536.9/18; PROEX 237/17).

In this study we conducted the experiment using a total of 56 nests, thus 112 couples of blue tits. In our experiment, only the first adult captured (25 males and 31 females) was temporarily manipulated by reducing the UV reflectance on yellow feathers. Plumage colour manipulation has no apparent negative effects in health or survival, as suggested by previous studies (e.g., Delhey et al. 2006, in adult blue tits). Video-recording inside the nest has no apparent effect on the behaviour or reproductive performance of wild birds, since parents resume offspring feeding normally in less than ten minutes and sometimes even in the presence of researchers. Also, in previous experiments in the study area, none of the experimental blue-tit nests was deserted during or after video-recording. Cross-fostering was performed between nests with similar laying date and clutch size. Manipulation of individuals lasted less than 5 minutes and did not involve blood sampling.

### Study area and study species

The study was carried out in Miraflores de la Sierra, Madrid, central Spain (40° 48’N, 03° 47’W) during the breeding season of 2018. We studied a wild population of blue tits (*Cyanistes caeruleus*) breeding in nests-boxes located in a deciduous forest, mainly dominated by Pyrenean oak (*Quercus pyrenaica*). In this species, nest construction and incubation is carried out by females, but both adults contribute to offspring provisioning, which is crucial for offspring growth and survival (Schwagmeyer & Mock 2008).

In our study area, blue tits raise one clutch per season. This species shows one of the largest clutch sizes in Passerines and may raise up to 15 nestlings (average brood size in the study population: 9.6 ± 1.8 SD; n = 464; range 4-15). Therefore, a blue tit family constitutes a complex social network, which connects a considerable number of family members that coincide for a short period of time and that interact to adjust their decisions over parental care. Thus, it is expected that signals of individual quality have evolved as a mechanism to facilitate information exchange. Indeed, both parents exhibit colourful plumage during the breeding season (Hunt et al. 1999). That is the case of the UV/blue colouration of crown feathers, which reflects condition (Delhey et al. 2006) and plays a role in mate choice (Andersson et al. 1998; Hunt et al. 1999). Also, UV/blue brightness of mates is related to rearing capacity, and male and female blue tits seem to follow opposite strategies according to this trait: when paired with a UV-reduced mate, females decrease their feeding rates, while males increase them (Limbourg et al. 2004; 2013b).

Both in offspring and adults, blue tits show a carotenoid-based colouration on yellow breast feathers, which also reflects light in the UV region (wavelength between 300-400 nm). Carotenoid colouration of yellow feathers is related both to genetic quality (Ferrer et al. 2015) and rearing capacity (Senar et al. 2002; García-Navas et al. 2012), and indicates health status both in offspring (Johnsen et al. 2003; Hidalgo-García 2006) and adults (Doutrelant et al. 2008; del Cerro et al. 2010). However, most of the evidence on yellow breast colouration working as a signal of quality is correlative and the function of UV reflectance of yellow plumage has been little explored (in nestling blue tits, see Johnsen et al. 2003; Jacot & Kempenaers 2007; Morales & Velando 2018; see also Galván et al. 2008 in the closely related great tit, *Parus major*). To our knowledge, UV colouration of yellow breast plumage has not been experimentally manipulated in adult birds and thus no study has explored whether this trait affects the behavioural strategies of parents and offspring, which is why we conducted the current study.

### General methods and experimental manipulation of UV reflectance

At the beginning of the breeding season, nest-boxes were visited every three days to register laying date, the onset of female incubation and hatching date (= day 0). Two days before the expected hatching date, all clutches were reciprocally cross-fostered between two nests, that were similar in clutch size and laying date (± 2 eggs and days respectively).

In the second week of the nestling period, we captured adults with nest-box traps. The first adult captured was assigned to either an experimental UV-blocked group, in which we reduced the UV/yellow reflectance of breast feathers, or to a Control group (n = 29 UV-blocked nests and 28 Control nests; hereafter UV-blocked individuals). In the UV-blocked treatment, we manipulated 13 males and 15 females from different nests by applying a yellow marker on breast feathers (Edding 4500, code 005), following previous studies in this study population (see Morales & Velando 2018; see also Galván et al. 2009, in great tits, *Parus major*). UV-blocked adults have lower UV reflectance, resembling low quality individuals. The adult that was captured first was also marked on the back feathers with a white permanent marker (Edding 751, code 049) in order to distinguish parents during video recording. At Control nests, we applied the same yellow marker but only on the inner tail feathers of the first adult captured (n= 12 males, n= 16 females; hereafter Control-treated individuals). This was done in order to block UV reflectance of a similar-sized region as in the experimental group (in case the marker has non-desirable side-effects that may obscure the observed behavioural patterns), but on an area that cannot be seen by potential receivers and thus cannot act as a signal. First adult captured in Control nests were also marked with white on the back. In all UV-blocked and Control nests, the second adult captured (i.e., the partner of the manipulated individual) did not receive any colour manipulation.

The original colour of yellow breast feathers was also measured using a portable spectrophotometer (JAZZ, Ocean Optics) and UV chroma was calculated as reflectance in the UV range divided by total reflectance (R_300–400_/R_300–700_; following Johnsen et al. 2005). Prior to treatment, UV-blocked and Control-treated individuals did not differ in UV/yellow chroma (F_1,54_ = 0.373; *P* = 0.54).

We captured adults until one day before the video recording, so that the manipulation could be perceived by all family members at least during one day (average number of days between plumage manipulation and video recording: 3.51 ± 0.71 SD days; range 1-4 days; n=56). On day 12 after hatching, all chicks were ringed, weighed, and individually marked at the head or wings with the same white marker used for adults. We also collected 3-5 breast feathers per chick for molecular sexing (see Supplementary material). On day 13, we video-recorded all nests to register the behaviour of all family members during the feeding bouts. Finally, all nests were visited days after to establish the number of nestlings that fledged (i.e., fledging success).

### Video recordings and behavioural analyses

On day 12, the original nest-box was substituted by a recording nest-box with a camera decoy so that the blue tit pair could become familiar with the set-up before recording. On day 13, we recorded the behaviour of all family members during parental provisioning by placing a night-vision video camera on the recording nest box (DX, 8 LED and 180° vision, China). The camera was placed on the opening at an approximate distance of 10 cm from the nest (Morales & Velando 2018). In total, we obtained the blue tit feeding behaviour of all nests except one Control nest. However, we excluded 14 nests in which we did not detect one of the adults during the whole video observation (the first adult captured did not appear in 3 nests and the second adult captured did not do so in 11). Thus, we ensured that all behavioural observations corresponded to nests with bi-parental care, since the behaviour of both manipulated individuals and their partners is needed to understand parental care strategies. The final sample size for behavioural observations was thus 43 nests (19 Control nests and 24 UV-blocked nests). We were unable to differentiate the first and the second adult in two nests, although they were clearly two different individuals, so for these two nests we could not analyse feeding variables for each parent separately, but we did include them in the models of total feeding rates. In one nest, both adults fed during the video recording but only one fed during the scored time. Thus, this nest was included in the analyses.

We recorded each nest during 1.5 hours but we observed 30 minutes per nest. The first 30 minutes were excluded to avoid possible disturbance due to researcher’s presence after leaving the camera on the nest. Video recordings were performed between 8:00 and 13:00, except 1 video that was recorded between 19:00 and 21:00, due to rainfall during the morning.

For each feeding event we registered i) whether the adult was marked or not, ii) the prey size it provided and iii) chick begging intensity. Prey size was rated on a 3-point scale: 1 = small (equal or shorter than the adult beak), 2 = medium (larger than the beak but smaller than the adult head), and 3 = large (bigger than the adult head) (see Morales & Velando 2018). To establish begging intensity, we rated it on a 4-point scale for each individual chick, following Kölliker et al. (1998): 0 = calm, 1 = weak gaping, 2 = gaping and neck stretched, 3 = gaping, neck stretched, and standing, 4 = gaping, neck stretched, standing, and wing flapping. All these variables were registered by an observer who was unaware of treatment. Once all behavioural variables were registered, the sum of feeding rates per nest and average prey size was obtained for both parents together and for each parent separately.

### Statistical Analyses

First, we analysed whether the number of feeding bouts differed between treatments. All parental behavioural variables (i.e., total number of feeding bouts and average prey size for both parents, and separately for manipulated individuals (either UV-blocked or Control-treated) and their partners) were analysed using General Linear Models (GLMs). In these models we included nest treatment (UV-blocked or Control), the sex of the manipulated adult, brood size at day 13 and days elapsed from treatment to video recording as predictor variables. Since treatment effects may differ between the sexes, we included the interaction between treatment and sex of the manipulated adult in all the models. In the models exploring the behaviour of each parent separately, we additionally included the corresponding behaviour (feeding bouts or prey size) of the mate as predictor variable.

Regarding chick individual begging intensity, we performed a mixed model that included nests ID as a random factor. In this model, we included treatment, sex of the manipulated adult, the interaction between treatment and sex of the manipulated adult, sex of the focal nestling, brood size and days after treatment. In the model of begging intensity directed to the manipulated parent (either UV-blocked or Control-treated), we controlled for begging intensity directed to the mate, and vice versa. Similarly, nestling body mass was analysed using a mixed model with nest ID as a random factor. In this model we included treatment, sex of manipulated adult, nestling sex, brood size, days after treatment and the interaction between treatment and sex of manipulated adult. In all mixed models, degrees of freedom were obtained using the Satterthwaite approximation. Fledging success was analysed using a Generalized Linear Model (GLZ) with Poisson distribution. In this model we included treatment, sex of manipulated adult, brood size, days after treatment and the interaction between treatment and sex of manipulated adult as covariates.

We used SAS 9.4 (SAS Inst., Cary, NC, USA) for all statistical analyses. When non-significant (α=0.05), the interaction between treatment and sex of the manipulated parent was removed. Models were checked for residual normality with a Shapiro test. All tests were conducted by using Type III sums of squares.

## Results

### Feeding bouts and prey size

Treatment significantly affected the total number of feeding bouts performed (Table 1), UV-blocked nests receiving overall more feeding bouts than Control nests (mean ± SE: 19.48 ± 1.66 and 15.63 ± 1.43, respectively). When analysing parental behaviour separately, there was a significant effect of treatment on the feeding behaviour of the partners: partners of UV-blocked individuals provided more feeding bouts than partners of Control-treated individuals (Table 1; Fig. 1). As expected, treatment did not affect the number of feeding bouts of manipulated parents, either UV-blocked or Control-treated (Table 1; Fig. 1). Days elapsed from manipulation to the video recording was negatively related with the total number of feeding bouts and with the number of the feeding bouts performed by the partner (Table 1). The sex of the manipulated adult did not affect the number of feeding bouts in any model (Table 1) and the interaction between treatment and sex of the manipulated adult was never significant (Table S1).

**Table 1.** General lineal models (GLMs) showing the effects of treatment on the number of feeding bouts performed by both parents and by each parent separately (i.e., the manipulated adult and the partner). Manipulated refers to UV-blocked individuals in UV-blocked nests and Control-treated ones in Control nests. The interaction between treatment and the sex of the manipulated parent was never significant (see Table S1). Coefficients are shown for Control nests and for females. Significant differences are marked in bold.

|  | ***Feeding bouts***<br>***of both***<br>***parents*** | ***Feeding bouts***<br>***of the partner*** | ***Feeding bouts of***<br>***manipulated***<br>***adult*** |
| --- | --- | --- | --- |
| ***Intercept*** | *coef* = 29.18 ± 8.99 | *coef* = 21.59 ± 7.40 | *coef* = 14.93 ± 8.20 |
| ***Treatment***<br>***(Control)*** | *coef* = -4.73 ± 2.13<br>*F*<sub>1,37</sub> = 4.91<br>***P* = 0.033** | *coef* = -3.81 ± 1.82<br>*F*<sub>1,35</sub> = 4.38<br>***P* = 0.044** | *coef* = -2.12 ± 1.90<br>*F*<sub>1,35</sub> = 1.25<br>*P* = 0.27 |
| ***Sex of***<br>***manipulated***<br>***adult***<br>***(females)*** | *coef* = -0.11 ± 2.07<br>*F*<sub>1,37</sub> = 0.00<br>*P* = 0.96 | *coef* = 0.44 ± 1.75<br>*F*<sub>1,35</sub> = 0.06<br>*P* = 0.80 | *coef* = 0.43 ± 1.75<br>*F*<sub>1,35</sub> = 0.06<br>*P* = 0.81 |
| ***Brood size*** | *coef* = 0.61 ± 0.59<br>*F*<sub>1,37</sub> = 1.06<br>*P* = 0.31 | *coef* = 0.21 ± 0.50<br>*F*<sub>1,35</sub> = 0.18<br>*P* = 0.67 | *coef* = 0.59 ± 0.49<br>*F*<sub>1,35</sub> = 1.45<br>*P* = 0.24 |
| ***Days after***<br>***treatment*** | *coef* = -4.13 ± 1.49<br>*F*<sub>1,37</sub> = 7.68<br>***P* = 0.0087** | *coef* = -3.01 ± 1.27<br>*F*<sub>1,35</sub> = 5.58<br>***P* = 0.024** | *coef* = -2.21 ± 1.31<br>*F*<sub>1,35</sub> = 2.82<br>*P* = 0.10 |
| ***Feeding bouts***<br>***provided by the***<br>***other parent*** |  | *coef* = -0.29 ± 0.16<br>*F*<sub>1,35</sub> = 3.11<br>*P* = 0.087 | *coef* = -0.29 ± 0.16<br>*F*<sub>1,35</sub> = 3.11<br>*P* = 0.086 |

**Figure 1.**
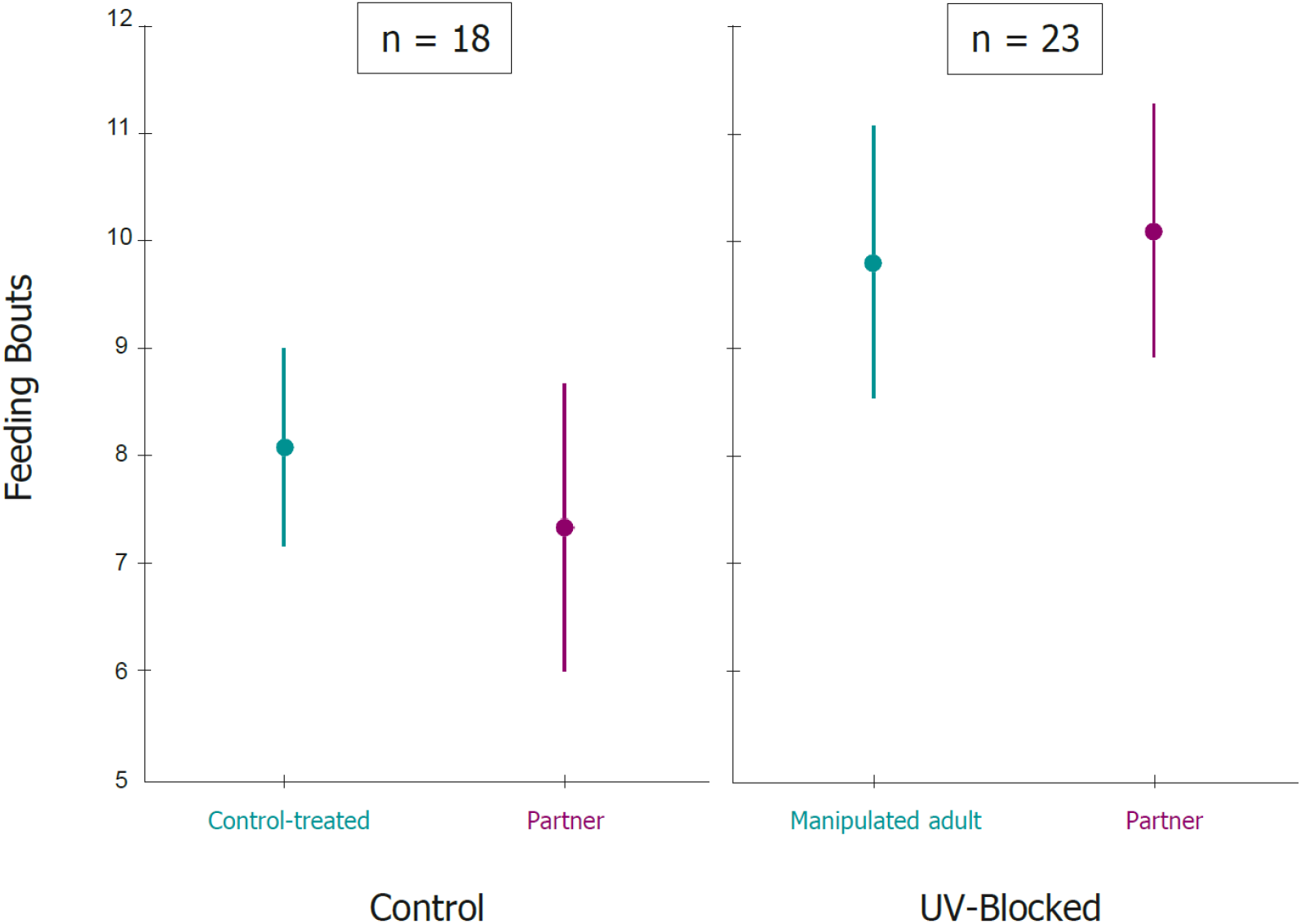
Number of feeding bouts performed by manipulated adults (UV-Blocked/Control-treated) and their partners. Error bars denote standard errors (mean ± SE; n=41). Sample sizes for each treatment are shown.

Parents of UV-blocked nests provided smaller prey items than parents of Control nests (mean ± SE: 2.22 ± 0.06 and 2.44 ± 0.07, respectively; Table 2). However, there was no treatment effect on prey size when analysed for each parent separately (Table 2), although the prey size of UV-blocked adults tended (non-significantly) to be lower than that of Control-treated adults (Table 2). The interaction between treatment and sex of the manipulated adult was not significant (see Table S2).

**Table 2.** General lineal models (GLMs) showing the effect of treatment on the prey size mean per nest provided by both parents and for each one separately (i.e., the manipulated adult and the partner). Manipulated refers to UV-blocked individuals in UV-blocked nests and Control-treated ones in Control nests. The interaction between treatment and the sex of the manipulated parent was never significant (see Table S2). Coefficients are shown for Control nests and for females. Significant differences are marked in bold.

|  | ***Prey size***<br>***provided by***<br>***both parents*** | ***Prey size***<br>***provided by***<br>***the partner*** | ***Prey size***<br>***provided by***<br>***manipulated***<br>***adult*** |
| --- | --- | --- | --- |
| ***Intercept*** | *coef* = 1.67 ± 0.39 | *coef* = 1.16 ± 0.65 | *coef* = 1.79 ± 0.55 |
| ***Treatment***<br>***(Control)*** | *coef* = 0.26 ± 0.09<br>*F*<sub>1,37</sub> = 7.57<br>***P* = 0.0091** | *coef* = 0.12 ± 0.15<br>*F*<sub>1,34</sub> = 0.64<br>*P* = 0.43 | *coef* = 0.24 ± 0.12<br>*F*<sub>1,34</sub> = 3.74<br>*P* = 0.062 |
| ***Sex of***<br>***manipulated adult***<br>***(females)*** | *coef* = -0.03 ± 0.09<br>*F*<sub>1,37</sub> = 0.11<br>*P* = 0.75 | *coef* = -0.11 ± 0.13<br>*F*<sub>1,34</sub> = 0.69<br>*P* = 0.41 | *coef* = -0.19 ± 0.12<br>*F*<sub>1,34</sub> = 2.56<br>*P* = 0.12 |
| ***Brood size*** | *coef* = 0.03 ± 0.03<br>*F*<sub>1,37</sub> = 1.73<br>*P* = 0.20 | *coef* = 0.05 ± 0.04<br>*F*<sub>1,34</sub> = 1.92<br>*P* = 0.18 | *coef* = -0.01 ± 0.03<br>*F*<sub>1,34</sub> = 0.13<br>*P* = 0.73 |
| ***Days after***<br>***treatment*** | *coef* = 0.07 ± 0.07<br>*F*<sub>1,37</sub> = 1.22<br>*P* = 0.28 | *coef* = 0.03 ± 0.09<br>*F*<sub>1,34</sub> = 0.09<br>*P* = 0.76 | *coef* = 0.07 ± 0.08<br>*F*<sub>1,34</sub> = 0.65<br>*P* = 0.43 |
| ***Prey size provided***<br>***by the other adult*** |  | *coef* = 0.24 ± 0.19<br>*F*<sub>1,34</sub> = 1.58<br>*P* = 0.22 | *coef* = 0.19 ± 0.15<br>*F*<sub>1,34</sub> = 1.58<br>*P* = 0.22 |

### Begging intensity and nestling fitness parameters

There was no treatment effect on the begging intensity of nestlings, neither to both parents (Table 3) nor when analysed for each parent separately (Table 3; Fig. 2). Begging intensity was lower in larger broods (Table 3). The interaction between treatment and sex of the manipulated adult was not significant (Table S3).

**Table 3.** General mixed models (GLMMs) showing the effect of treatment on the mean begging intensity of nestlings directed to both parents and to each one separately (i.e., the manipulated adult and the partner). Manipulated refers to UV-blocked individuals in UV-blocked nests and Control-treated ones in Control nests. The interaction between treatment and the sex of the manipulated parent was never significant (see Table S3). Coefficients are shown for Control nests and for females. Significant differences are marked in bold.

|  | ***Begging intensity to both parents*** | ***Begging intensity to the partner*** | ***Begging intensity to manipulated adult*** |
| --- | --- | --- | --- |
| ***Intercept*** | *coef* = 1.31 ± 0.58 | *coef* = 0.96 ± 0.43 | *coef* = 0.19 ± 0.48 |
| ***Treatment (Control)*** | *coef* = 0.11 ± 0.14<br>*F*<sub>1,36.5</sub> = 0.67<br>*P* = 0.42 | *coef* = 0.01 ± 0.11<br>*F*<sub>1,33.4</sub> = 0.01<br>*P* = 0.92 | *coef* = 0.11 ± 0.11<br>*F*<sub>1,34.8</sub> = 1.00<br>*P* = 0.33 |
| ***Sex of manipulated adult (females)*** | *coef* = -0.06 ± 0.13<br>*F*<sub>1,36.2</sub> = 0.22<br>*P* = 0.64 | *coef* = -0.18 ± 0.10<br>*F*<sub>1,32.4</sub> = 3.27<br>*P* = 0.080 | *coef* = -0.18 ± 0.11<br>*F*<sub>1,34.1</sub> = 2.73<br>*P* = 0.11 |
| ***Sex of nestling (females)*** | *coef* = -0.02 ± 0.06<br>*F*<sub>1,326</sub> = 0.15<br>*P* = 0.70 | *coef* = -0.01 ± 0.07<br>*F*<sub>1,321</sub> = 0.05<br>*P* = 0.83 | *coef* = -0.01 ± 0.06<br>*F*<sub>1,316</sub> = 0.01<br>*P* = 0.93 |
| ***Brood size*** | *coef* = -0.09 ± 0.04<br>*F*<sub>1,38.9</sub> = 4.90<br>***P* = 0.033** | *coef* = -0.07 ± 0.03<br>*F*<sub>1,37.5</sub> = 4.94<br>***P* = 0.032** | *coef* = -0.007 ± 0.03<br>*F*<sub>1,39.1</sub> = 0.04<br>*P* = 0.84 |
| ***Days after treatment*** | *coef* = 0.13 ± 0.09<br>*F*<sub>1,35.5</sub> = 1.94<br>*P* = 0.17 | *coef* = 0.03 ± 0.07<br>*F*<sub>1,32.4</sub> = 0.21<br>*P* = 0.65 | *coef* = 0.10 ± 0.08<br>*F*<sub>1,33.9</sub> = 1.85<br>*P* = 0.18 |
| ***Begging intensity to the other parent*** |  | *coef* = 0.64 ± 0.05<br>*F*<sub>1,316</sub> = 184.15<br>***P* < 0.0001** | *coef* = 0.55 ± 0.04<br>*F*<sub>1,332</sub> = 179.19<br>***P* < 0.0001** |

**Figure 2.**
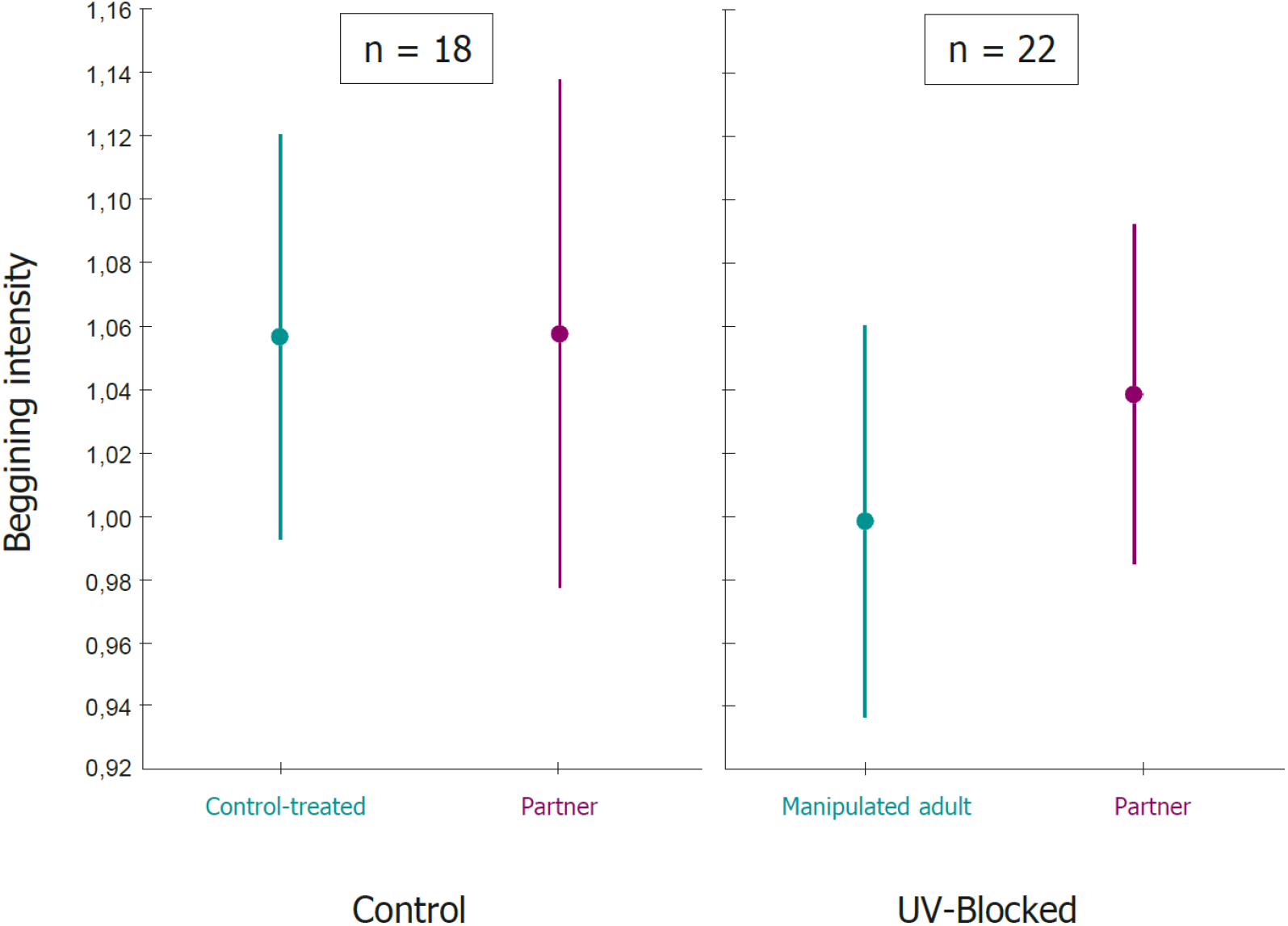
Begging intensity mean of nestlings to manipulated adults (UV-Blocked/Control-treated) and their partners. Error bars denote standard errors (mean ± SE; n=40). Sample sizes for each treatment are shown.

No effect of treatment was found on nestling body mass (Table 4). Male nestlings were significantly heavier than females (Table 4). Brood size was negatively related to nestling body mass (Table 4). Finally, no effect of treatment was found on the number of fledglings (Table 4). The interaction between treatment and sex of the manipulated adult was not significant (Table S4).

**Table 4.** General mixed model (GLMM) and Generalized lineal model (GLZ) with Poisson distribution showing the effects of treatment, respectively, on nestling body mass fledging success. Coefficients are shown for Control nests and for females. The interaction between treatment and the sex of the manipulated parent was never significant (see Table S4). Significant differences are marked in bold.

|  | ***Nestling Body mass***<br>***(g)*** | ***Fledging success*** |
| --- | --- | --- |
| ***Intercept*** | *coef* = 11.55 ± 0.53 | *coef* = 0.96 ± 0.48 |
| ***Treatment***<br>***(Control)*** | *coef* = -0.04 ± 0.14<br>*F*<sub>1,43.2</sub> = 0.10<br>*P* = 0.76 | *coef* = 0.02 ± 0.12<br>*χ*<sup>2</sup><sub>1</sub> = 0.03<br>*P* = 0.87 |
| ***Sex of manipulated***<br>***adult***<br>***(females)*** | *coef* = 0.01 ± 0.13<br>*F*<sub>1,42.7</sub> = 0.00<br>*P* = 0.96 | *coef* = -0.02 ± 0.11<br>*χ*<sup>2</sup><sub>1</sub> = 0.02<br>*P* = 0.88 |
| ***Sex of nestling***<br>***(females)*** | *coef* = -0.44 ± 0.06<br>*F*<sub>1,362</sub> = 65.64<br>***P* < 0.0001** |  |
| ***Brood size*** | *coef* = -0.15 ± 0.04<br>*F*<sub>1,44.8</sub> = 14.73<br>***P* = 0.0004** | *coef* = 0.13 ± 0.03<br>*χ*<sup>2</sup><sub>1</sub> = 17.02<br>***P* < 0.0001** |
| ***Days after***<br>***treatment*** | *coef* = 0.06 ± 0.08<br>*F*<sub>1,42</sub> = 0.49<br>*P* = 0.49 | *coef* = -0.0004 ± 0.08<br>*χ*<sup>2</sup><sub>1</sub> = 0.00<br>*P* = 0.99 |

## Discussion

In this experimental study we investigated whether UV colouration may act as a condition-dependent signal of parental quality, which may have (co-)evolve in multiple contexts (Morales & Velando 2018). UV colouration could be perceived by nestlings and affect their begging behaviour (i.e., the amount of food they demand), and by the mate, who may co-adjust its contribution to parental care accordingly. To test this hypothesis, we experimentally blocked the UV colouration of yellow breast feathers of one blue tit adult per nest. We indeed found that blue tits coupled with UV-blocked mates increased the number of feeding bouts, providing experimental evidence that UV colouration acts as a signal that affects parental care strategies in blue tits. However, despite of this, we found no evidence that nestlings responded to this parental signal. Below we discuss the potential causes and consequences of this discrepancy.

### Parental behaviour: Feeding bouts and prey size

By blocking the UV reflectance on yellow breast feathers, we mimicked the appearance of low-quality individuals (Johnsen et al. 2003; Jacot & Kempeanaers 2007). We found that the parental investment, here measured in terms of the total number of feeding bouts – thus considering both parents – was significantly higher in nests with an UV-blocked adult. When we analysed the pair members separately, we found that the observed increase was due to the fact that partners of UV-blocked adults increased the number of feeding bouts. We hypothesize that the reduction in UV reflectance of yellow plumage triggered a compensatory response by the partner, who took a greater proportion of care given the apparent lower-quality partner, as found in previous studies in this and other species (Gowaty et al. 2003: Bluhm & Gowaty 2004; Bolund et al. 2009; Limbourg et al. 2013b). Interestingly, we found that both males and females responded similarly to changes in mate’s UV/yellow reflectance, suggesting similar rules over investment in response to this trait. Previous studies on the same species showed that males and females responded differently when paired with non-attractive mates with experimentally reduced UV/blue crown colouration: blue tit females decreased their feeding rates when paired with UV-blocked males (Limbourg et al. 2004), whereas males increased their feeding rates when paired with UV-blocked females (Limbourg et al. 2013b). The causes of this differences remain as yet unresolved. Although UV/blue colouration has been widely studied in blue tits (Hunt et al. 1999; Limbourg et al. 2004; 2013b; 2013b; Lucass et al. 2016), few studies have explored the signalling function of UV/yellow colouration of breast feathers. This study is the first evidence that UV colouration of breast feathers also acts as a signal of quality in adults during the reproductive period.

UV-blocked adults did not change their rate of nestling provisioning. Indeed, the condition or quality of the UV-blocked adult had not changed after manipulation but only the signal expression. The manipulated adult is only supposed to change its behaviour if it self-perceives the manipulated signal directly or if it adjusts its behaviour because its responds indirectly to the changes in its partner’s behaviour (Burley 1986; 1988; Sanz 2001; Cline et al. 2016), for which we did not have evidence. The latter is in contrast to a recent hypothesis that partners co-adjust their investment based on behavioural traits - by checking each other during feeding visits, which should result in an alternated sequence of feeding visits (Hinde et al. 2005; Johnstone et al. 2014, Iserbyt et al. 2019; Griffioen et al. 2019). The fact that our manipulation did induce a unilateral change in parental provisioning rather suggests that partners respond to the mate’s signal, and not to its behaviour. However, it is also possible that partners responded to both mate’s cues combined. Even though this remains speculative, we propose that partners could rate their mate’s parental effort differently after manipulation, as they expect a lower contribution of care by their mate - based on the (manipulated) signal. Thus, the partner perceives that the UV-blocked adult is performing relatively better than expected given its (manipulated) low quality. In other words, as its signal but not its feeding rate has changed, this could trigger a positive response in the partner.

Finally, UV colouration of yellow feathers may signal rearing capacity rather than attractiveness, otherwise we should have found a decrease in partner’s feeding effort, because a reduction in the perceived quality of the partner lowers the current value of the brood and thus the willingness to invest (Burley 1986; 1988; Kokko 1998; Ratikainen & Kokko 2010). Our results also show that in nests with an UV-blocked adult both parents provided prey items smaller than Control nests. One possibility is that the increase of total feeding rates in UV-blocked nests would compromise the provisioning over larger prey items. However, this would only apply to the parents that increased feeding rates (i.e., the partners of UV-blocked individuals). A non-exclusive possibility is that UV-blocked adults relaxed their foraging effort due to the increase of investment of their partners.

### Offspring: Begging, growth and survival

As a blue tit family consist of a complex social network in which signals could be perceived by all family members, we expected that the UV manipulation in adults could affect parent-offspring interactions. Indeed, parental signals have been seen to affect offspring behaviour, although evidence is scarce (but see Tinbergen & Perdeck 1950; Velando et al. 2013), and blue tit nestlings have been shown to respond to their siblings’ UV/yellow colouration (Morales & Velando 2018), revealing that the signal can be perceived. If UV colouration of breast feathers signals low rearing capacity, we hypothesised that offspring could perceive UV-blocked parents as poor-quality caregivers. Thus, chicks should beg less intensively to manipulated parents due to expected low returns, and assuming that begging is energetically costly (Chappell et al. 2002). However, we did not find differences in nestling behaviour in function of the experimentally manipulated parental trait. This could be due to the fact that offspring – contrary to adults – do not respond to the signal but to other cues, such as the number of feeding bouts, prey-testings or to adult’s feeding calls. We argue that the signal might not be relevant for the offspring, who may only have to distinguish between parents, and then associate the respective phenotype with a given parental quality. However, adults may use this signal before the rearing period when direct information on adult provisioning is not available yet.

We did not find effects of our treatment on nestling body mass nor on the number of fledglings. One possible explanation is that our manipulation was too late, so that nestlings are already nearly fully developed and growth differences more difficult to detect. Nonetheless, the signal may still be relevant even at later stages, since parent-offspring bonds continue for weeks after fledging until full independence (Salvador 2016; Stenning 2018). Another possible explanation could be that the time between manipulation and mass measurement was too short (on average 3.51 days ± 0.71 SD; see Material & Methods). Nonetheless, this seems less likely given that, previously, the blocking of UV/yellow nestling colour resulted in changes in brood behaviour and body mass after 24 h (Morales & Velando 2018). Finally, in UV-blocked nests, parents increased their feeding bouts but reduced prey size, so overall the energetic intake might have been equal, causing the lack of differences between treatments in chick body mass.

### Conclusions

Our study shows that UV colouration of yellow breast feathers in adult blue tits affects parental feeding strategies. Through UV-blocking manipulation, we observed that adults responded to the partner’s manipulated trait and not to its behaviour (i.e., feeding bouts) in order to adjust their parental effort. This supports the concept of flexible parenting, with UV/yellow colouration of blue tit adults revealing aspects of parental capacity and parents adjusting their investment based on this signal. The opposite effect was found for offspring: nestlings did not respond to the manipulated signal of one of their parents, but we hypothesize that they should rather respond to their parents’ behaviour (feeding bouts, prey-testings or feeding calls). However, too little attention has thus far been paid to how offspring could optimally fine-tune their behaviour to parental rearing capacity.

## Notes

### Competing Interest Statement

The authors have declared no competing interest.

